# Exo-Tox: Identifying Exotoxins from secreted bacterial proteins

**DOI:** 10.1101/2025.04.04.647182

**Authors:** Tanja Krueger, Damla A. Durmaz, Luisa F. Jimenez-Soto

**Author notes:** Contributing authors.

## Abstract

**Background:** Bacterial exotoxins are secreted proteins able to affect target cells, and associated with diseases. Their accurate identification can enhance drug discovery and ensure the safety of bacteria-based medical applications. However, current toxin predictors prioritize broad coverage by mixing toxins from multiple biological kingdoms and diverse control sets. This general approach has proven sub-optimal for identifying niche toxins, such as bacterial exotoxins. Recent Protein Language Models offer an opportunity to improve toxin prediction by capturing global sequence context and biochemical properties from protein sequences.

**Results:** We introduce Exo-Tox, a specialized predictor trained exclusively on curated datasets of bacterial exotoxins and secreted non-toxic bacterial proteins, represented as embeddings by Protein Language Models. Compared to Basic Local Alignment Search Tool (BLAST)-based methods and generalized toxin predictors, Exo-Tox outperforms across multiple metrics, achieving an Matthews correlation coefficient > 0.9. Notably, Exo-Tox’s performance remains robust regardless of protein length or the presence of signal peptides. We analyze its limited transfer-ability to bacteriophage proteins and non-secreted proteins.

**Conclusion:** Exo-Tox reliably identifies bacterial exotoxins, filling a niche overlooked by generalized predictors. Our findings highlight the importance of domain-specific training data and emphasize that specialized predictors are necessary for accurate classification. We provide open access to the model, training data, and usage guidelines via the LMU Munich Open Data repository.

## 2 Introduction

Bacterial toxins include the most potent toxins known today. Many have been directly linked to deadly human diseases, such as botulinum toxin, cholera toxin, or diphteria toxin. It has been hypothesized that bacteria produce them for defense (e.g. evasion of predators) or to allow the colonization of preoccupied habitats Speare et al (2018); Sana et al (2016); Carbonetti et al (2007). Whatever their objective is, their origins are diverse, and many are known to originate from horizontal gene transfer Kumar et al (2019); Brouwer et al (2020) through bacteriophages.

When considering bacterial toxins, it is necessary to distinguish between endotoxins and exotoxins. The first are of lipidic nature and released upon physical disruption of bacterial membranes. The second, exotoxins, are actively secreted proteins. Their secretion occurs either into the extracellular space or directly into the target cells involving diverse sets of pathways and secretion signals Rietschel et al (1996); D’Onofrio and Paradisi (1983); Sheehan et al (2022). Our focus is on bacterial exotoxins, which we define as: secreted proteins capable of cellular manipulation of a target organism.

With the identification of bacterial toxins, specifically diphteria toxin in the 1800s Roux and Yersin (1888), a new understanding of human diseases caused by bacteria started, leading to the development of a vaccine against diphteria, even though the connection had not always been immediately understood.

With emerging bacteria based treatments like probiotics Suez et al (2019); Lerner and Matthias (2020); Merenstein et al (2023) or fecal microbiota transplantation Holvoet et al (2016); Malard et al (2021), any method able to predict bacterial exotoxins from protein sequences will improve their safety. Identification of exotoxins may also lead to the discovery of novel drug candidates, considering that some exotoxins have been proven to be suitable pharmaceutical agents Choudhury et al (2021); Poulain et al (2015); Peigneur and Tytgat (2018).

Traditional wet-lab approaches like genomewide mutagenesis have helped to identify toxins and their importance in infection for bacteria growing in the lab. For larger scale, this approach is not economically feasible or possible. *In-silico* methods using the amino acid sequence of proteins are a promising alternative. Several of these approaches include techniques based on sequence similarity analyzing and clustering tools, (e.g.BLAST Altschul et al (1990), PSI-BLAST Altschul et al (1997); or MMseqs2 Steinegger and Söding (2017)), and homology methods based on Hidden Markov Models(HMMs) Krogh et al (1994) such as HMMER Eddy (2011). These, however, are often limited to implicit (HMMs) or explicit (BLAST) local sequence motives to identify toxins Alley et al (2020).

Efforts to predict toxins using Machine Learning vary in their focus and approach. Many models are designed specifically Pan et al (2019); Naamati et al (2009); Cole et al (2019); Gacesa et al (2016), or peptides Gupta et al (2013); Wei et al (2021, 2022). By contrast, other predictors Saha and Raghava (2007); Ahn et al (2022); de Nies et al (2021) expand their scope to include a broader set of virulence factors - proteins associated with pathogenicity, and are not limited to classical toxins. More recently, generalized toxin predictors have emerged combining toxins from multiple biological origins, including animals, plants, and bacteria Gupta et al (2013); Caceres-Delpiano et al (2021); Morozov et al (2023). Predictors often included very diverse non-toxin control sets, including proteins from different origin species than the toxins, or proteins with sub-cellular localizations distinct from secreted toxins Cole et al (2019); Caceres-Delpiano et al (2021); Morozov et al (2023).

Toxins predictors also differ in their input features and architecture. The most recent, CSM-Toxin Morozov et al (2023), uses a modification of ProteinBERT Brandes et al (2022), a protein language Model (pLM), and aims to find any type of toxin independent of origin. pLMs are pre-trained models that learn contextual relationships between amino acids in a protein sequence, analogous to how language models capture the meaning of words within a sentence. After training on large protein databases, the pLM encodes sequences into numerical vector called embeddings. These numerical representations can then serve as input for specialized downstream models.

Taking advantage of the T5 architectures used for the pLM model ProtT5 Elnaggar et al (2022), and realizing the lack of a bacterial exotoxin predictor trained in only bacterial exotoxins, we present here Exo-Tox: a specialized predictor that closes the gap of identifying bacterial exotoxins from secreted non-toxic bacterial proteins using the protein’s primary sequence and protein Language Models (pLMs). Our model uses a highly curated dataset of bacterial exotoxins, defined here as secreted proteins capable of cellular manipulation of a target organism. As negative label, non-toxic secreted bacterial proteins were used. Exo-Tox is the name given to our best performing model, after evaluation of two predictors with different sets of input features: a naive approach of amino acid composition (*aac*), and protein embeddings generated by protT5 (Embs20). Exo-Tox identifies the toxin potential of secreted bacterial proteins more reliably than a generalized toxin prediction tool, a BLAST approach, or CSM-Toxin predictor. We investigated the relevance of signal peptides, and protein length in their performance. None of them play a role in the capacity of the predictor to classify toxins. To evaluate the applicability and transferability of the Exo-Tox, we applied it to two related proteins sets, bacteriophages (natural transmitters of toxins) and general bacterial proteins, and we present these results.

## 3 Methods

### 3.1 General information

We built a predictor to differentiate bacterial exotoxins and secreted bacterial non-toxin. We compared two input feature approaches to a sequence similarity based method. Input features included the amino acid composition and an embedding based feature set. For the selected approach we investigated the biological relevance and robustness twofold: by testing partial sequences without the signal peptides and by retraining on scrambled sequences.

### 3.2 Data accessibility

All code used for data analysis is accessible in the LMU University repository https://doi.org/10.5282/ubm/data.576. As stated in the data availability statement, raw data and code for data wrangling that support the findings of this study are openly available in LMU repository.

### 3.3 Raw Data

The raw data consists of a set of expert curated bacterial exotoxins and a set of secreted, bacterial non-toxins previously published https://doi.org/10.5282/ubm/data.423 and analyzed under Krueger et al (2024). The dataset contains exotoxins from four bacterial toxin types, its sequences downloaded from Swiss-Prot (UniProtKB/Swiss-Prot (RRID:SCR 021164)) Bairoch and Apweiler (1996) and PubMed (PubMed (RRID:SCR 004846)) Sayers et al (2022). The database includes only active subunits of toxins. All sequence labeled as “fragment” or “partial” were removed. We also excluded any other virulence factors, and protein-based toxins that were not produced by bacteria. The total number of exotoxins is 2396 sequences. The non-toxins set is based on the PSORTb 3.0b, a database containing the predicted sub-cellular localization of bacterial protein.Yu et al (2010). Sequences with predicted association to membranes, outer membrane vesicles (OMVs), periplasm, or cytoplasm were removed, leaving only secreted proteins(9082 sequences). The full description of the data curation is described in Kruger et al, 2024 Krueger et al (2024).

### 3.4 Redundancy reduction

Raw datasets may contain sequences that are highly similar or identical to each other, as a result from biological processes (e.g horizontal gene transfer), or from artificial sources (e.g. sampling bias during data collection, labeling). To avoid overestimating the predictive power of our models by including redundant sequences, we performed two steps of redundancy reduction using the MMseqs2 algorithm Steinegger and Söding (2017). First, we removed duplicates between toxins and non-toxins with MMseqs2 using the easy-cluster option and its default parameters. We manually set the similarity threshold of 100% and the alignment coverage mode to 0, which causes that both the query and the target sequences must be fully covered by the alignment at the specified threshold. We retained only the representative sequence from each cluster, and removed any sequences that were present in both the toxin and non-toxin sets from the non-toxin control proteins. Second, we reduced the sequence similarity within the toxins and non-toxins datasets independently with a similarity threshold of 30%. This step was carried out before the separation of the test set and the cross validation folds. This approach makes sure that no two proteins between the sets shared more than the selected 30% sequence similarity. The resulting redundancy-reduced datasets contain 1069 toxins and 1308 non-toxins.

### 3.5 Generating the test set

We generated a hold-out test set, through stratified splitting, with the label as stratifying factor and a split ratio of 85% training/validation and 15% test sequences. The selection of the test set did not use other datasets (benchmarks) because of their inclusion of proteins that were non-toxins but virulence associated, or from different kindoms. Considering we reuse this test-set to compare our work with the existing predictor CSM-Toxin, our final test set was equally redundancy reduced against our own- and the CSM-Toxin training set. For this we used MMseqs2 with a similarity threshold of 30% and modus 0. Any subsequent steps of scaling, or features selection was performed independently on the training and validation set, without taking the test set into account.

### 3.6 Vector representation of protein sequences - Input features

#### 3.6.1 Generation of vectors of amino acid composition

We tried different protein representations as input features. For the naive approach, we used the amino acid composition (*aac*) of proteins calculated with equation 1.The resulting feature vector contains 20 individual values, which were scaled as described under “Prediction Method” section.

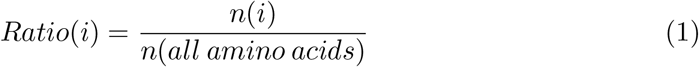

where:

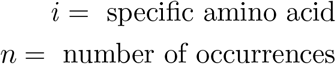

#### 3.6.2 Generation of embeddings based input features

To obtain the embedding representation of each protein, we used the pre-trained Protein Language de(pLM) ProtT5 Elnaggar et al (2022) version ProtT5-XL-UniRef50 (also referred to as ProtT5-XL-U50). We used their embed ProtT5.ipynb colab notebook to translate the protein sequences to per-protein embeddings which consists of 1024 values (https://colab.research.google.com/drive/1TUj-ayG3WO52n5N50S7KH9vtt6zRkdmj?usp=sharing#scrollTo=tRe7CfuqFFmY, accessed June 2023). To prevent overfitting, we applied a Principle Component Analysis (PCA) and retained the first 20 principal components from the 1024 embedding dimensions. (See “Prediction Method”) These 20 principal components based on embeddings (Embs20) are considered a separate approach to input features to the *aac* features for subsequent training.

### 3.7 Prediction method

We applied several supervised machine learning algorithms to predict toxins. The use of embeddings outsources computationally expensive pre-training, allowing us the use of computationally less demanding algorithms such K-Nearest Neighbors (kNN), Logistic Regression (LR), Support Vector Classifier (SVC). Random Forest (RF) and extreme Gradient Boosting (XGB). We scaled the input features for kNN, LR, and SVC using a Standard Scaler. No scaling was performed for RF and XGB, as these models are not affected by the feature scale. We then reduced the dimensionality of the embeddings by PCA to the first 20 Principal Components. To prevent data-leakage between the training, validation and test set, we used and a pipeline to carry out all steps. The pipeline fits the mentioned pre-processing steps to each fold of the training set and then performs the transformations o the validation and test data. Model hyper-parameters were optimized via Gridsearch on the training set. Table 2-6 in the SOM summarize which hyper-parameters were searched and the respective search spaces. The search space was set up on a logarithmic scale around the default values as described in the sklearn documentation (1.5.1). Any hyper-parameters not specifically mentioned were set to default. The hyper-parameter performance was measured by 10 fold cross validation using Matthews Correlation Coefficient (MCC) as balanced performance metric Chicco et al (2021). The architecture’s hyperparameter combination with the highest cross-validation score was selected. The performance on an unseen dataset was measured using a holdout test set, which was kept out of feature transformation and parameter optimization. No further model changes were made on the basis of the test set performance. For details on program and package versions see Supplementary file *environment*.*yml* in repository.

### 3.8 BLAST

As a second baseline, we found the closest match in sequence similarity using Blastp in the command line tool from NCBI National Center for Biotechnology Information (2008). We first constructed a custom BLAST database out of the training sequences and then ran the established test set sequences as query, choosing the query with the lowest e-value. The labels from resulting matches were compared to the ground truth labels of the test sets.

### 3.9 Performance measures

We evaluated the model performance on a variety of metrics. We followed the common practice of labeling true positives as (TP), false positives (FP), true negatives (TN) and false negatives as (FN). TP are proteins that are correctly predicted as toxic. FP are the proteins that are wrongly predicted as toxins. TN are proteins that are correctly predicted as non-toxic, and FN are toxins not classified as such.

#### 3.9.1 Performance metrics during model optimization

As previously mentioned, we used the Matthew’s Correlation Coefficient (MCC) to measure model performance during hyper-parameter optimization and model selection. MCC was picked for its balanced penalty for FP and FN classified observations. MCC was calculated using equation 2.

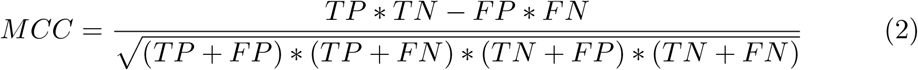

where:

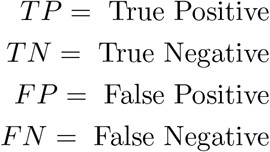

#### 3.9.2 Performance metric of final predictor

For better comparability with other predictors additional performance metrics to the MCC were calculated on the finished predictors using the hold-out test set. They include accuracy (Equ. 3), precision (Equ. 4), recall (Equ. 5), ROC-AUC. Bootstrapping was applied to receive a more reliable metric that takes outliers in the data into account. Each metric was averaged across 10000 samples that were chosen at random with replacement. The equations for each metric are listed below with N being the number of bootstrapping samples, and b for the index of each bootstrap-sample.

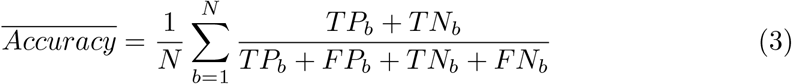

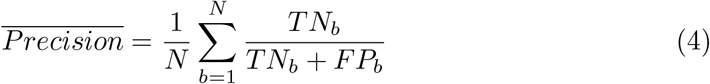

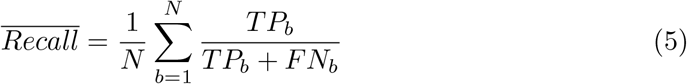

To get the 95% confidence interval for each metric, the Standard Error *SE*_*metric*_ was multiplied by 1,96.

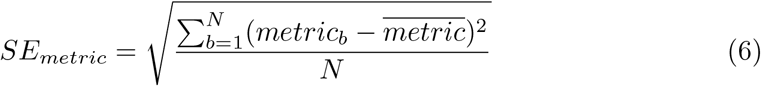

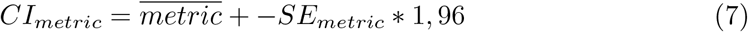

*CI*_*metric*_ = *metric* + −*SE*_*metric*_ * 1, 96 (7) With *CI*_*metric*_ as the confidence interval of a metric, N being the number of bootstrapping samples, *metric*_*b*_ being the metric of choice for a single bootstrap sample b and 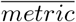 as the mean across all bootstrap samples.

### 3.10 Investigating the robustness of the models

#### 3.10.2 Evaluation of Signal Peptide bias

Both raw datasets of toxins and non-toxin consist of secreted proteins. We therefore investigated the impact of signal peptides on the performance of our predictor. For this, the SignalP-6.0 Teufel et al (2022) was used on both datasets, setting the model mode to “fast”, and the organism option to prokaryotes (“other”). We removed the predicted signal peptides from the original amino acid sequences in our test set. We then applied the same methodology outlined in Generation of embeddings based input features and Generation of vectors of amino acid composition to the truncated test sequences.

#### 3.10.2 Evaluation of length bias

To investigate the impact of sequence length, we scrambled all sequences and obtained new embeddings using ProtT5. For the scrambling, an out-of-bag sampling with no replacement was done creating a new artificial sequence. For each sequence we generated embeddings, and used them to retrain the best performing model using Support Vector Classifier and the first 20 Principal Components of a PCA. Through shuffling, the new embeddings are restricted to information of sequence length and amino acid composition. To evaluate the effect of length information on training with embeddings, we compared this re-trained model to our baseline (*aac*)-which only uses amino acid composition as input and contains no length information.

#### 3.11 Predictor scope - Generation of additional datasets

Some bacterial toxins are known to originate from bacteriophages. We therefore tested Exo-Tox on a dataset of bacteriophage proteins. Our bacteriophage set combines four previously published phage datasets. This includes: the PhaNNs database Cantu et al (2020), phage accession IDs from the EMBL Database (European Nucleotide Archive (ENA) (RRID:SCR 006515), European Bioinformatics Institute (RRID:SCR 004727)), the full Actinobacteriophage Dataset Russell and Hatfull (2017) and a dataset from Zhang et al Zhang et al (2015). We removed all proteins labeled as fragment, partial sequences or prophage. Prophages were removed, because their sequence is affected by the evolution of the organism’s DNA in which they are embedded, which can change the uniqueness as toxin. We further reduced the set to 30 percent sequence similarity.

To investigate the limits of the specialized predictor Exo-Tox, which was trained on secreted proteins only, we tested the model on other bacterial proteins. For this we chose all proteins from the reference proteomes as published under NCBI: https://www.ncbi.nlm.nih.gov/datasets/genome/ (Accessed in Jan, 2024) from the same species IDs that were found for the bacterial toxins. Any sequences that were part of our exotoxin set were removed. Again, we removed any fragments or partial sequence. In addition any sequences that were already identical and reduced the redundancy to 30% sequence similarity.

To investigate the possibility of unidentified toxins in the control set, we performed an extensive search using regular expressions applied to the protein descriptions provided by the fasta files. Regular expressions were used to identify proteins with descriptions indicative of toxin-like activity. The list of regex terms used in this analysis is detailed in the appendix (see S7 and S8)

### 3.12 Use of Generative AI

Each section was first researched, drafted and written by the authors. Sections of the manuscript were then passed to language models including DeepL Write, ChatGPT version 4o, and Anthropic version Heiku to correct grammar, revise code, restructure arguments for better clarity, and checking spelling mistakes.

## 4 Results

### 4.1 Embeddings distinguish exotoxins from non-toxins

The relative percentage of each amino acid and their order, defines the physico-chemical properties and functions of proteins. In previous work, we have shown that secreted bacterial proteins and exotoxins have different amino acid usage Krueger et al (2024). However, amino acid composition (aac) cannot capture the global context between amino acids. Protein embeddings created by Protein Language Models (pLM) such as ProtT5, are designed to capture this global context. To explore if aac and protein embeddings distinguish between bacterial exotoxins and bacterial secreted non-toxins, we applied a Principle Component Analysis (PCA) to the data. The first two principal components are visualized in 2D plots. (Fig 1.)

**Fig. 1:**
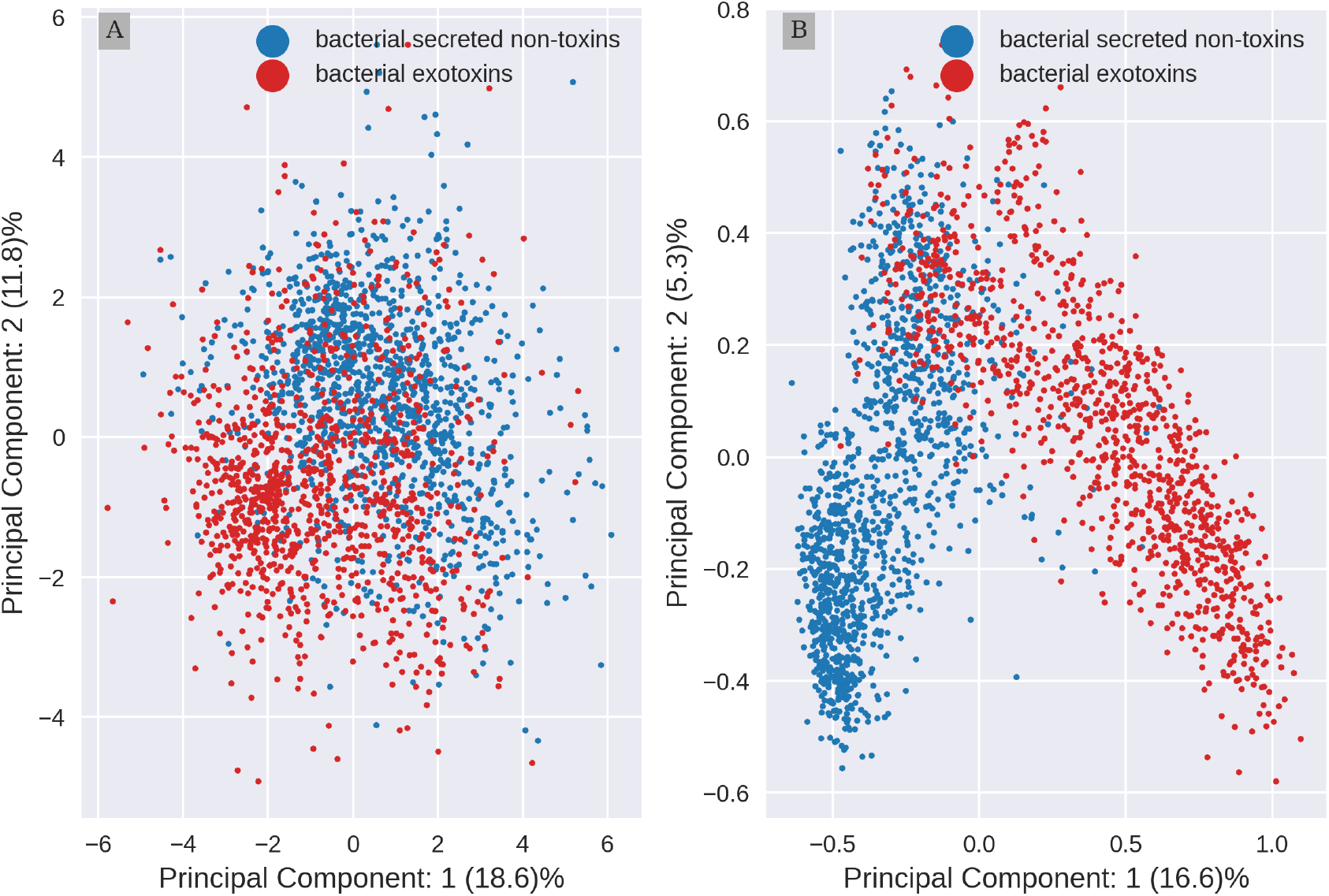
Embeddings information separates better toxins from non-toxins than aac. A) 2D Principal Component Analysis(PCA) projection of amino acid composition (aac) (B) and per protein embeddings by ProtT5. The aac was scaled before the PCA was carried out, embeddings were not scaled (see material and methods). Bacterial exotoxins (red) and secreted bacterial non-toxins (blue) form overlapping clusters.

This visual analysis shows both representations of proteins form clusters, detecting differences between toxins and secreted non-toxic proteins (control). For aac, PCA (Fig 1A) uncovers two disperse clusters with extensive overlap along the whole range of the second principal component (PC 2). In contrast, for embeddings (Fig 1B) the clusters are more compact, with a small overlap limited to the upper half of the PC 2. These results indicate that representation of secreted proteins and toxins in embeddings capture more information than aac, possibly allowing for a better separation of the two classes of proteins when creating a predictor model.

### 4.2 Embeddings based prediction outperforms aac

Both representations detect differences between secreted and toxic proteins. Therefore, we evaluated if the information contained in them is enough to predict toxicity potential of secreted proteins by training a series of simple supervised models. We compared two approaches of input features. A naive approach using the aac as input, and a second method using dimensionality reduced embeddings. (Embs20). As baseline, we used BLAST alignment as classification method, which does not rely on machine learning, but rather finds motifs of sequence similarity between proteins.

Five machine learning architectures were tested. They included K-Nearest Neighbors (k-NN), Support Vector Classifier (SVC), Logistic Regression (LR), Random Forest (RF) and extreme Gradient Boosting (XGB). In all five, Embs20 outperformed *aac* as input feature. (Fig 2). Differences based on the type of input used, are statistical relevant when evaluating the 95% confidence interval(CI). In contrast, differences in architecture are negligible, as interpreted by the overlapping of CI ranges for both inputs sets for kNN, SVC, LR, RF and XGB.

**Fig. 2:**
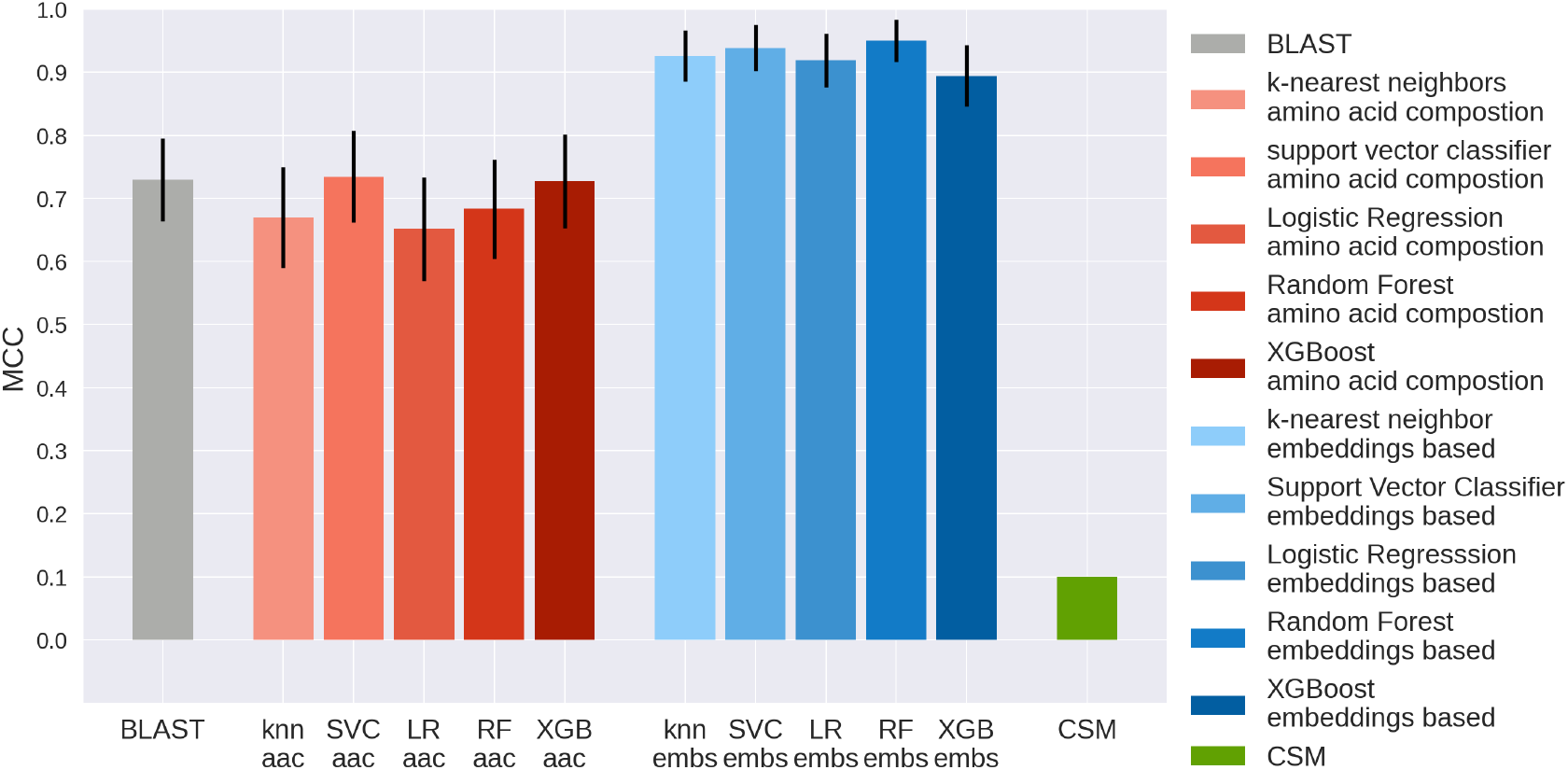
Toxicity of secreted proteins predicted accurately from embeddings. Different exotoxins prediction methods are compared. This includes two sets of input features. Methods that use the amino acid composition as input are depicted in shades of red. Methods using the first 20 Principle Components that retain the most information form ProtT5 protein embeddings are in shades of blue. The two input features approaches are compared to BLAST as a baseline (gray) and CSM-Toxin, a state of the art generalized toxin predictor not specialized on bacterial proteins (green)Morozov et al (2023). Data: hold-out test set of bacterial exotoxins and bacterial, secreted non-toxins with less then 30% sequences similarity to training set of CSM-Toxin and training set of our proposed predictor. Metric: Matthew’s correlation coefficient (MCC). Model architectures:kNN: K-Nearest Neighbors, LR: Logistic Regression, SVC: Support Vector Classifier, RF: Random Forest and XGB: extreme Gradient Boosting. The model architectures are differentiated by color intensity.From light to dark: kNN,LR,SVC,RF and XGB. Black whiskers mark the 95% interval with +-the 1.96 the standard error. Embeddings based approach paired out-performed acc baseline, the generalized predictor method and BLAST. The different model architectures perform similarly.

All metrics for the evaluation of all methods are in table 1. BLAST performed similar to the *aac* baseline, with MCC of 0.73 and accuracy of 0.86. For precision and recall, however, the performance ranking changed. BLAST recall of 0.71 is below the naive aac method with values between 0.78 and 0.87. This suggest that BLAST has a higher probability to miss true toxins.

**Table 1:** Embeddings-based predictor outperformed other approaches.

| Method | MCC | Accuracy | Precision | Recall | ROC AUC |
| --- | --- | --- | --- | --- | --- |
| BLAST | 0.731± 0.066 | 0.858± 0.038 | 0.972± 0.031 | 0.708± 0.074 | - |
| **Amino Acid Composition** |  |  |  |  |  |
| kNN | 0.670± 0.080 | 0.837± 0.040 | 0.836± 0.061 | 0.796± 0.064 | 0.919± 0.029 |
| SVC | 0.735± 0.073 | 0.868± 0.036 | 0.843± 0.057 | 0.871± 0.054 | 0.950± 0.020 |
| LR | 0.652± 0.082 | 0.828± 0.040 | 0.828± 0.062 | 0.783± 0.066 | 0.906± 0.034 |
| RF | 0.683± 0.079 | 0.843± 0.039 | 0.830± 0.060 | 0.824± 0.061 | 0.931± 0.026 |
| XGB | 0.727± 0.075 | 0.865± 0.037 | 0.851± 0.058 | 0.851± 0.058 | 0.942± 0.023 |
| **Embs20** |  |  |  |  |  |
| kNN | 0.926± 0.040 | 0.963± 0.020 | 0.941± 0.037 | 0.980± 0.023 | 0.986± 0.012 |
| SVC | 0.938± 0.036 | 0.969± 0.019 | 0.948± 0.035 | 0.986± 0.019 | 0.995± 0.006 |
| LR | 0.919± 0.043 | 0.960± 0.021 | 0.953± 0.034 | 0.959± 0.032 | 0.994± 0.005 |
| RF | 0.950± 0.034 | 0.975± 0.017 | 0.973± 0.026 | 0.973± 0.026 | 0.992± 0.007 |
| XGB | 0.895± 0.049 | 0.948± 0.024 | 0.933± 0.040 | 0.953± 0.034 | 0.981± 0.014 |
| CSM-Toxin | 0.140± 0.097 | 0.571± 0.054 | 0.785± 0.223 | 0.075± 0.042 | - |

Matthew Correlation Coefficient (MCC), Accuracy, Precision, Recall, and Receiver Operating Characteristic - Area Under the Curve (ROC-AUC) for different exotoxin prediction approaches. Methods include BLAST, the CSM-Toxins predictor Morozov et al \(2023\), and two different input feature approaches paired with five machine learning architectures. Input features are the amino acid composition (aac) and the first 20 Principal Components of a Principal Component Analysis performed on T5 per-protein embeddings (Embs20). Architectures include kNN (K-Nearest Neighbors), LR (Logistic Regression), SVC (Support Vector Classifier), RF (Random Forest), and XGB (Extreme Gradient Boosting). The models paired with Embs20 input features show the best performance across all metrics.

**Table 2:** Protein length does not influence prediction.

|  | MCC | Accuracy | Precision | Recall | ROC AUC |
| --- | --- | --- | --- | --- | --- |
| aac/SVC |  |  |  |  |  |
| original sequences | 0.735± 0.073 | 0.868± 0.036 | 0.843± 0.057 | 0.871± 0.054 | 0.950± 0.020 |
| Embs20/SVC |  |  |  |  |  |
| scrambled sequences | 0.685± 0.078 | 0.840± 0.040 | 0.788± 0.062 | 0.885± 0.052 | 0.908± 0.031 |

Matthew Correlation Coefficient (MCC), Accuracy, Precision, Recall, and Receiver Operator Curve - Area Under the Curve (ROC-AUC) for two different inputs. Methods include the Embs20/SVC trained on randomly scrambled sequences and the aac/SVC predictor that was introduced in table 1.

As benchmark for the predictor, we used CSM-Toxin Morozov et al (2023). This is a state of the art general toxin predictor, trained on a set including animal and bacterial toxins. Using our test set in the CSM-Toxin predictor gave a surprisingly low MCC score (0.14).The low value is reflected by the CSM predictions, where it correctly identified 174 out of 177 non-toxins, but missed the majority of the bacterial toxins by identifying 11 out of 147.

### 4.3 Signal peptides do not influence prediction

All sequences in this study are secreted proteins, but not all toxins have traditional signal sequences. We investigated which secretion signals are present in the data using

SignalP-6.0. Signal Peptides distinguish between two translocation routs: Sec and Tat and three Signal Peptidases: SPI-III. SignalP includes SP: Sec/SPI, LIPO: Sec/SPII, TAT: Tat/SPI, LIPOTAT: Tat/SPII, PILIN: Sec/SPIII and OTHER indicates no known signal peptides. We found that the presence of signal peptides differs between the toxins and non-toxins. For over 90% of the toxins no known secretion signal was identified by SignalP-6.0, while only 27% of the non-toxins contained no predicted secretion signal (see figure 3, Panel A). The most frequent signal peptides were the sec signals SP (378) and LIPO (470) occurrences in the control proteins. Because of this imbalance, and the fact that context is captured by the embeddings, we investigated if signal sequences could be introducing a bias in the Embs20 predictor.

**Fig. 3:**
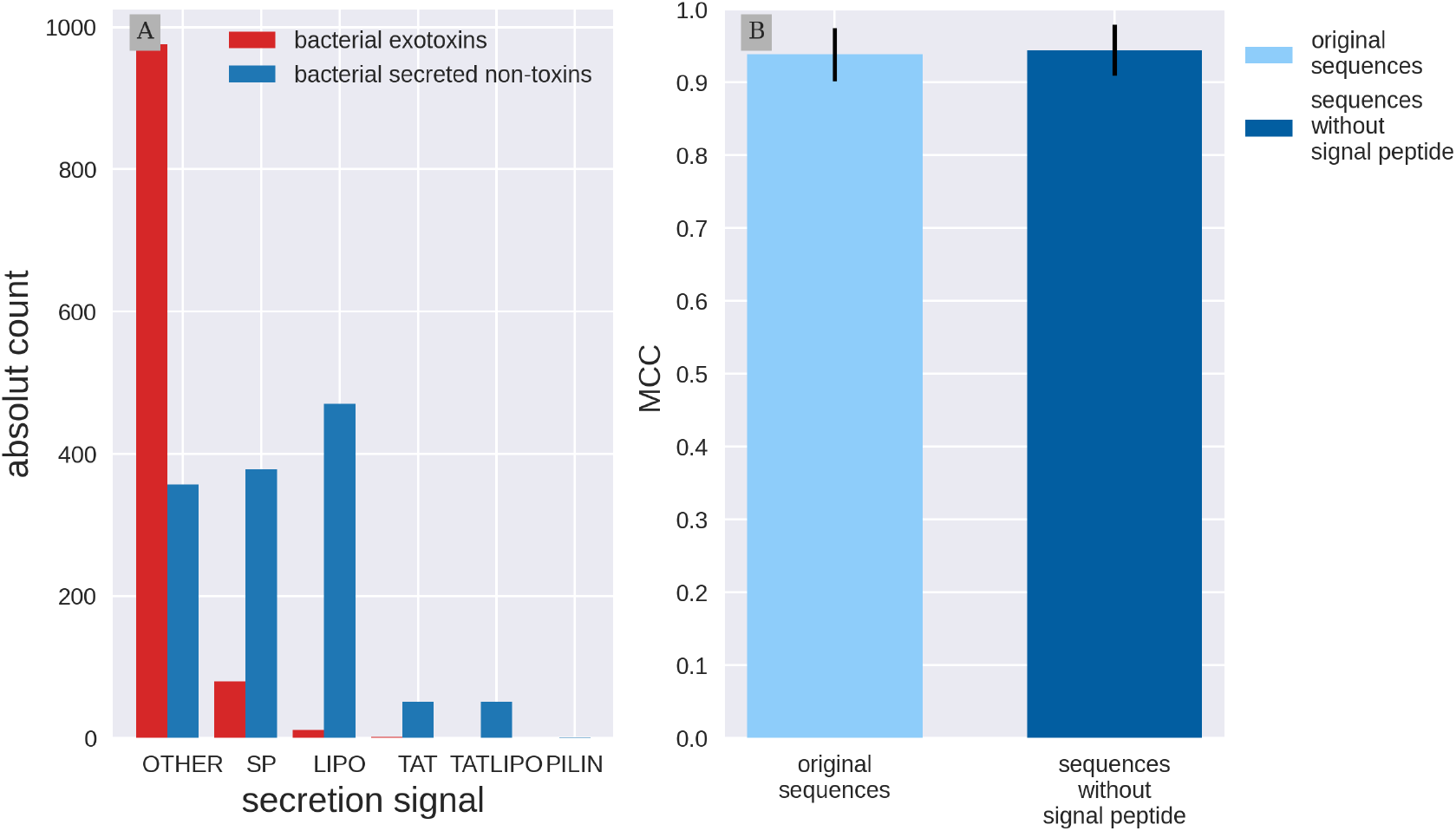
Signal Peptides do not influence toxin prediction. Panel A: Exploration of signal peptides predicted with SignalP-6.0. Bacterial exotoxins are in red, the bacterial secreted non-toxins are in blue. Signal Peptides distinguish between two translocation routs Sec and Tat and three Signal Peptidases SPI-III. Prediction include SP: Sec/SPI, LIPO: Sec/SPII, TAT: Tat/SPI, LIPOTAT: Tat/SPII, PILIN: Sec/SPIII and OTHER indicates no known signal peptides. The majority of exotoxins does not have a predicted signal peptide. Panel B: Performance comparison with and without signal Peptides. Model architecture was introduced in Fig 2 Support Vector Classifier (SVC) using the first 20 Principal Components calculated on per protein protT5 embeddings (Embs20). Two versions of the test set are compared. Light blue are the original test set sequences. Dark blue: the test sequences without the predicted signal peptides. Embs20/SVC performs equally well on both test set versions.

Therefore, we removed the predicted signal peptides from the test set sequences and obtained new embeddings to evaluate the performance once again. We compared the predictor performance on the original and modified test sets using the Embs20/SVC model architecture(Fig 3B). Embs20/SVC showed similar results on both sets, with no significant differences in the measured metrics, as the MCC shows (for both sets the MCC is around 0.94). This suggests that the signal peptide has little influence on the information captured by the embeddings and, therefore, has no effect in the predictors ability to identify toxicity in secreted bacterial proteins. Additional metrics are listed in table S1.

### 4.4 Protein length does not influence prediction

Previously, we identified differing median sequence lengths between our two sets Krueger et al (2024), raising concerns about potential bias. To examine the impact of these length variations, we retrained our best performing model (Embs20/SVC) on scrambled sequences. Their embeddings should reflect protein length and amino acid composition, but a different context. We compared this retrained model to our baseline which solely captures the amino acid composition, but not the protein length. (see baseline in fig 2). Both methods without context information yield comparable performance across all metrics, including an MCC of around 0.7 (see table 2). The consistent results across accuracy, precision, recall, and ROC-AUC suggest that the protein length doesn’t skew exotoxins predictions in the Embs20/SVC model, allowing us to assume that the information included in the embeddings used by the Embs20 model is not influenced by protein length.

### 4.5 Predictor limitations

We know that the embedding-trained predictor (Emb20/SVC) has been trained with context information, and that length and signal peptides information have not introduced a bias in the embedding-based selection process. The next step is to investigate the limitations of this model and it applicability to a new group of proteins. For this, we tested Embs20/SVC on bacteriophages, and on a set of bacterial proteins independent of their secretion status. (For details on the preprocessing of both sets, see Materials and Methods).Bacteriophages were chosen as many lysogenic phages contain toxin genes (prophages). The predictor classifies 159637 bacteriophage proteins as toxins (97%) and 54903 (63%) of bacterial proteins as toxins. (See Table 3). For control of performance, we also tested our full dataset of training and testing combined.

**Table 3:** Datasets and predictor results.

| Dataset | Toxin (predicted) | Non-toxin (predicted) |
| --- | --- | --- |
| **Embs20/SVC** |  |  |
| phages (reduced) | 159637 | 3787 |
| bacterial control (reduced) | 54903 | 31273 |
| original exotoxins | 2360 | 36 |
| secreted non toxins | 67 | 9015 |
| **aac/SVC** |  |  |
| phages (reduced) | 127344 | 36080 |
| bacterial control (reduced) | 54316 | 31860 |
| original exotoxins | 2150 | 246 |
| secreted non toxins | 700 | 8382 |

Predictions obtained for exotoxins and secreted non-toxins using Embs20/SVC (embedding-based) and aac/SVC (amino acid composition-based) classifiers across different datasets.

In this case, 2360 (98%) of the toxin data set is correctly classified as toxins, while from the secreted bacteria proteins only 67 (0.007%) were wrongly classified as toxins.

For comparison, the same analysis was performed using our best naive model based on amino acid composition (*aac*/SVC). In this case, 127344 (78%) of phage proteins are classified as toxins, and from the set of general bacterial proteins, 54316 (63%) were classified as toxins, which are similar to the results yielded from Embs20/SVC. The classification of exotoxins and secreted bacterial proteins set gave the expected results with 2150 (90%) of the toxins, and only 700 (8%) of secreted proteins identified as toxins. Although the toxin and secreted proteins sets are mostly correctly classified, results with bacteriophage proteins and bacterial proteins are unexpected.

### 4.6 Common classification by both predictors

Because of the previous results of the predictor on bacteriophage and general bacterial set, we examined which proteins were classified as toxins by both, the amino acid based (*aac*) and the embeddings-based (Embs20) predictor. The Upset plot allows us to visualize the different results and the number of common proteins classified by the predictors (Fig 4). The *Set Size* reflects the differences or similarities in number of proteins classified as toxins. The *Intersection Size* allows to see the numerical value of proteins in the evaluated set or intersection of sets, the dots indicate which set is being evaluated, and the lines connecting different sets show between which sets the proteins intersect. From all datasets, 125764 phage proteins are classified as toxins by both models (78% of Embs20/SVC, 98,7% of aac/SVC), and 39817 control bacterial proteins (“controls”) (78% of the Embs20/SVC toxins, 73% of the aac/SVC toxins). While 1276 exotoxins were classified as toxins by both models (54% of the Embs20/SVC, 59% of the aac/SVC), none of the proteins from the secreted dataset were identified as toxins by both models.

**Fig. 4:**
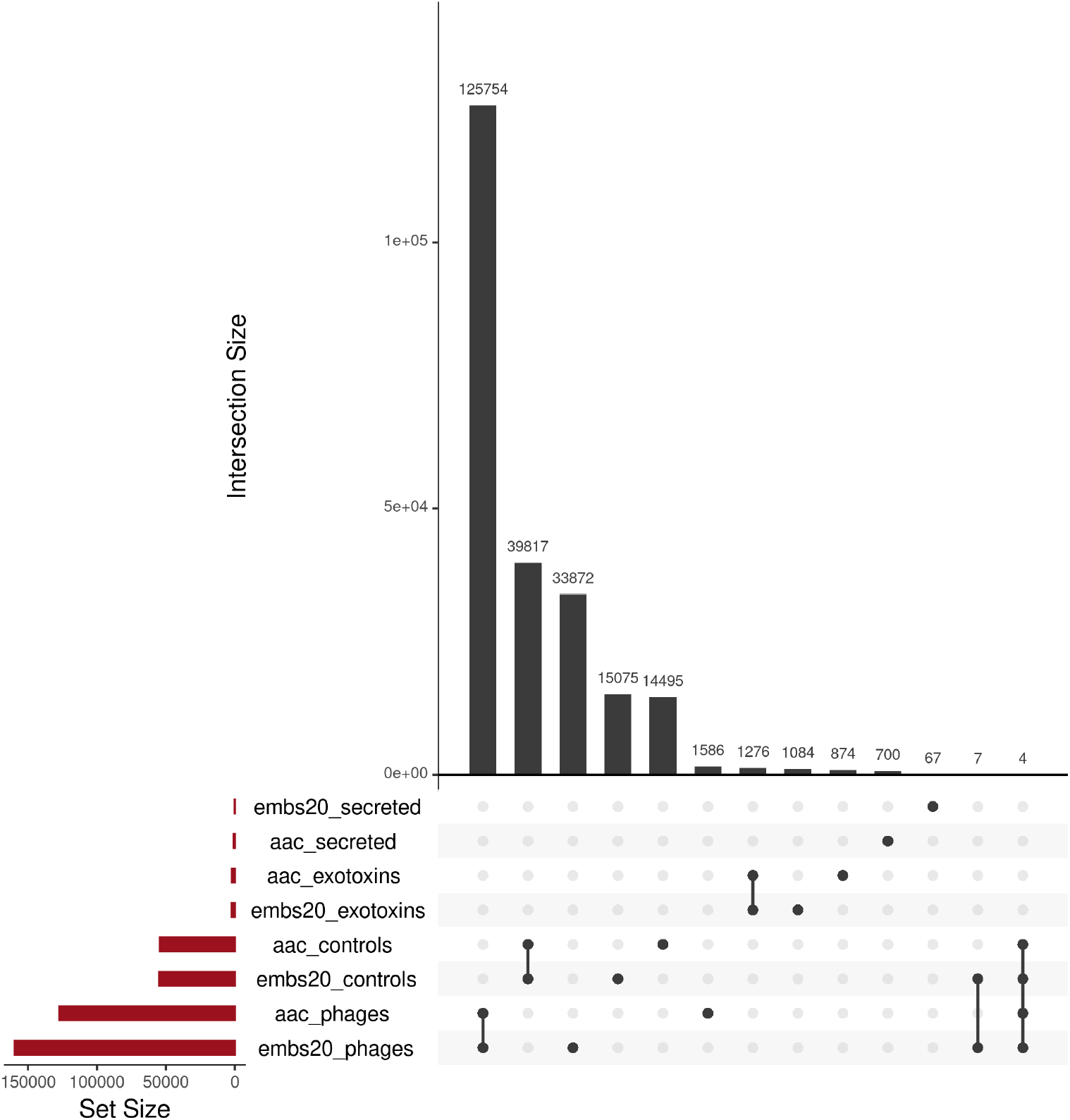
Several proteins are identified as toxins by both predictors, but not all. Size of the sets is shown in the lower left corner. The intersection of each set’s prediction are shown by the dots and bars located below each of the bars on the bar plot above. Four sets were tested 1) secreted: secreted bacterial proteins, 2) exotoxins: bacterial exotoxins, 3) controls: bacterial proteins independent of secretion status and 4) phages: phage proteins not containing prophages. Two predictors were tested 1) aac: was trained on amino acid composition and 2) Embs20: was trained on the first 20 Principle Components of per-protein embeddings.

## 5 Discussion

Although previously many toxins classifiers have been created, their development has morphed from highly specialized to very broad. Specialized predictors focused on specific biological kingdoms Naamati et al (2009); Cole et al (2019); Gacesa et al (2016), defined protein length Naamati et al (2009); Wei et al (2022); Gupta et al (2013), or were associated with distinct physiological activities Saha S (2007). Newer models are often trained on multi-species data sets Morozov et al (2023); Sharma et al (2022); Datta et al (2020) or exclusively on animal toxins Zhao et al (2022). While their wide focus may not always be explicitly stated in the literature, it can be inferred from the absence of constraints tied to specific use-cases. Some of these broader models have shown poor performance for domain-specific challenges, such as bacterial exotoxins Pan et al (2020).

To close this gap, we created predictors specialized in the classification of secreted bacterial exotoxins. The best performing model, Embs20/SVC, was renamed Exo-Tox. Unlike other predictors, Exo-Tox avoids the inclusion of virulence factors and endotoxins. It also narrows its training scope to secreted bacterial proteins to enhance performance within this specific domain. By selecting proteins from the same kingdom and sub-cellular localization, we aimed at minimizing extraneous features that separate the toxins and controls.

Considering the recent application of Natural Language methods for the creation of protein Language Models, Exo-Tox uses ML algorithms used for protein representation by embeddings using global context information that can be found in the sequence of proteins Elnaggar et al (2022). With Matthews Correlation Coefficient (MCC) of above 0.9, Exo-Tox provides a reliable tool in distinguishing secreted bacterial exotoxins from secreted non-toxins. This embedding based approach outperformed both the naive method of amino acid composition, and BLAST, (Table 1) implying that more information about protein toxicity is encoded in the global amino acid sequence (embeddings) than is found by local sequence similarity (BLAST) or in the simple amino acid composition. In performance metrics MCC, accuracy, recall, and ROC-AUC the embedding-based method outperformed the BLAST baseline. However, its precision was on par with the BLAST-based approach. Its ranking in precision and recall indicates that the model offers unique advantages: In the case of identifying novel toxins, high precision is critical to minimize the time of validating many positive hits in the lab, saving time and materials. For this application both the embedding based method and BLAST are equally reliable. In medical settings, however, where bacterial protein based interventions are screened for toxicity, a high recall rate becomes paramount to ensure no toxins go undetected. In this setting Exo-Tox is best suited for this task.

CSM-Toxin Morozov et al (2023) and Exo-Tox models use context information of protein sequence encoded by pre-train protein Language Models, The challenges of generalizing predictors were made visible with our test set: CSM-Toxin is trained on 90% eukaryotic and 10% prokaryotic origin, and in agreement with their own results in their test set, it achieves only a MCC of 0.14 on our test set of bacterial exotoxins. These results reflect the challenges of cross-kingdom predictive capacity, given the model’s eukaryotic-centric training data. This aligns with previous findings by Pan *et al*., who reported significant performance drops (from MCC 0.793 to 0.022) when animal-toxin models were tested on bacterial toxins Pan et al (2020). Their conclusion agrees with our own prior observations where we report similar distinctions between animal and bacterial toxins in terms of length, isoelectric point, and sequence similarity Krueger et al (2024). These results emphasize the importance of domain-specific training data.

We investigated potential biases by signal peptides (SPs) or protein length, and found no significant difference between Exo-Tox and comparative architecture predictors failing to contain this two aspects. SPs are crucial for certain secretion pathways; however, using SignalP, we observed that most toxins lacked identifiable SPs, suggesting alternative secretion systems (like T1SS, T3SS, T4SS, or T6SS).When we assessed bias from SP presence, we truncated SPs from primary sequences and retrained the predictor. Both predictors (with and without SP information) performed equally, confirming that SPs do not bias the training. Additionally, despite prior analysis identifying significant differences in median sequence lengths between groups Krueger et al (2024), predictors trained with and without length information performed comparably. This demonstrates that neither SPs nor protein length significantly impact model performance, validating our selection of secreted bacterial proteins as an appropriate negative set.

We tested the transferability of knowledge of Exo-Tox on bacteriophages and bacterial proteins. They were chosen given the relation of bacterial exotoxins with bacteriophages and their usage by bacteria. Bacteriophages are considered the origin of many bacterial exotoxins Naureen et al (2020); Jamet et al (2017); Boyd and Brüssow (2002). Exo-Tox classified 97% of bacteriophages proteins as toxins. This is unlikely to be accurate. Current research does not suggest that every phage protein is a toxin. This over-classification can lie in a dataset bias: toxins in our training data disproportionately originate from phages, leading Exo-Tox to associate phage-related patterns with toxicity. In the set of bacterial proteins, including non-secreted ones, Exo-Tox classifed about 60% as toxins. Investigating which proteins were classified both by amino acid composition and which by embeddings, revealed a large overlaps. This indicates that aac alone has a large influence.

A recent publication addresses the poor transferability of the model to biological molecules. They consider that the splitting of the training and test sets is the key for transferability Ektefaie et al (2024). Although our predictor’s training uses a combination of the randomness and sequence similarity reduction, which are the factors named by the authors, it will be interesting in the future to use their platform for splitting of the sets to determine if this plays a role. As for now we can only hypothesize that i) The embeddings principal components chosen for training are capturing bacteriophage specific information, or ii) the small data set for toxins contain a bias we have not been able to control causing an overfitting of Exo-Tox. Our results show that length and signal sequence are not playing a role in the classification. Whatever the reason, these findings reveal a significant limitation: Exo-Tox is most reliable when used exclusively for distinguishing secreted exotoxins from secreted non-toxins, the domain for which it was trained.

While Exo-Tox demonstrates strong performance in its intended domain, expanding its applicability to all bacterial proteins remains a challenge. One solution will be to expand the training dataset. However, this can be only realized when more experimental backed sequences are available and their labels confirmed. Our efforts to expand our controls are summarized in the appendix (see regex tables S7 and S8), which revealed many potential false negatives. A predictor trained on such data would struggle to achieve high recall for true toxins, as the signal from mislabeled non-toxins could overwhelm the true positive signals. However, removing such entries without them being known toxins, might remove the most interesting edge cases, introducing a bias entirely dependent on the initial data curation. For a truly general predictor that can effectively extrapolate toxicity regardless of origin, training data would need to encompass all biological kingdoms. This would need to include plants, fungi, .Moshiri et al (2016); Soliman et al (2021) even bacteriophages. Yet, the practicality of this approach is hindered by the overwhelming abundance of unconfirmed animal and bacterial toxins in existing protein databases. In light of these considerations, we advocate for the development of more specialized predictors, by narrowing the focus to specific applications or biological domains, which should reflect the training data.

In general, Exo-Tox is a specialized predictor for secreted bacterial exotoxins, addressing a key gap left by generalized predictors with cross-kingdom training data. Exo-Tox demonstrates strong predictive performance, outperforming amino acid composition models, BLAST-based approaches, and generalized predictors like CSM-Toxin. We show that potential confounding factors, such as protein length and signal peptides, do not bias the models predictions. While the over-classification of bacteriophage proteins and bacterial proteins independent of secretion status highlight the impact of dataset bias, it also underscores the importance of domain-specific training data and the need for clear application scopes. To support reproducibility and replicability, we provide open access to the model, training data, and usage instructions via the Open Data repository of the LMU Munich: https://doi.org/10.5282/ubm/data.576

## Abbreviations

2D: two dimensional
aac: Amino Acid Composition
BLAST: Basic Local Alignment Search Tool
CI: Confidence Interval, here typically used as the 95% CI implying an interval between 1.96*Standard Error
CSM-Toxin: particular toxin predictor Morozov et al (2023)
Embs20: first twenty principle components calculated on embeddings using a PCA
embeddings: fixed-size vectors derived from pre-trained pLMs
Exo-Tox: selected high performance predictor introduced in this work called Embs20/SVC during training
FN: False Negative
FP: False Positive
HMMER: tool for finding sequence homologs and alignments Eddy (2011)
HMMs: Hidden Markov Models
pI: isoelectric point
kNN: K-Nearest Neighbors
LIPO: signal peptide indicating the combination of Sec/SPII
LIPOTAT: signal peptide indicating the combination Tat/SPII
LR: Logistic Regression
MCC: Matthews Correlation Coefficient
MMseqs2: Many-against-Many sequence searching tool for clustering sequences Steinegger and Söding (2017)
OMVs: Outer Membrane Vesicles
pLMs: protein Language Model
PC: principle components from a PCA
PCA: principle component analysis
PILIN: signal peptide indicating the combination of Sec/SPIII
ProtT5: particular pLM Elnaggar et al (2022))
PSI-BLAST: Position-Specific Iterative Basic Local Alignment Search Tool Altschul et al (1997)
PSORTb: 3.0b, subcellular localization prediction tool Yu et al (2010)
RF: Random Forest
ROC-AUC: Receiver Operator Curve - Area Under the Curve
SignalP-6.0: tool to predict the presence of signal peptides Teufel et al (2022)
Sec: Secretory pathway, a protein transport system
SE: Standard Error
SP: Signal Peptide
SPI-III: signal peptidases 1-3
SVC: Support Vector Classifier
T5: architecture choice in pLMs
T1SS: Type I Secretion System
T3SS: Type III Secretion System
T4SS: Type IV Secretion System
T6SS: Type VI Secretion System
TAT: signal peptide indicating the combination of Tat/SPI
Tat: twin arginine translocation system, a protein transport system
TN: True Negative
TP: True Positive
XGB: extreme Gradient Boosting

## 6 Declarations

- Funding This research was made possible by funding from BMBF under the grant number SSTDBB-16DKWN136B to LFJS.
- Competing interests The authors declare that they have no competing interests.
- Ethics approval and consent to participate Not applicable.
- Consent for publication Not applicable.
- Data availability The datasets supporting the conclusions of this article are available in the repository, https://doi.org/10.5282/ubm/data.576.
- Code availability All code is written in python or R. All code, corresponding documentation and information for non-programmers on how to reproduce the project is available under, https://doi.org/10.5282/ubm/data.576, too.
- Authors’ contribution TK: Did exploratory data analysis,pre-processed and trained the Machine Learning algorithm, analyzed and visualized the model performance of baseline and benchmark predictors, did the bias analysis, curated bacterial data and prepared the original draft. DD: Curated the bacteriophage data. LJ: Conceptualized the project, curated exotoxins data, visualized model performance, acquired the funding, did the project administration and supervision, organized the resources and edited the manuscript. All authors read and approved the final manuscript.
- Acknowledgments We would like to express our sincere gratitude to Burkhard Rost for his insightful discussions and for fostering an environment that facilitated this research. We also extend our thanks to his lab members at TUM Bioinformatics chair Sebastian Franz, Tobias Olenyi, and Duc Anh Le for their invaluable advice and help in this work. Jutta Schreier (LMU) and Josefine Lakatos for their assistance with administrative aspects of this project; to Celine Marquet (TUM) for her valuable contributions for the grant; to Michael Heinzinger (TUM) for his help with the T5 model; to Thomas Gudermann (LMU) for his continuing support of this project. We would also extend our thanks to the scientific community for contributing their experimental findings to public databases; and to all individuals and curators responsible for maintaining these indispensable repositories. Special appreciation goes to Henrik Nielsen and his team at DTH in Lyngby, Denmark, for their long-standing commitment to upkeeping the invaluable SignalP resources (RRID:SCR 015644).

## Appendix A

Signal Peptides Influence on Model Performance

**Table S1:** Signal Peptides Do Not Influence the Model Performance. *No signal peptides* represent the values obtained with the model trained on embeddings of sequences without the signal sequence region predicted by SignalP6.0.

|  | MCC | Accuracy | Precision | Recall | ROC AUC |
| --- | --- | --- | --- | --- | --- |
| Original Sequences | 0.938± 0.036 | 0.969± 0.019 | 0.948± 0.035 | 0.986± 0.019 | 0.995± 0.006 |
| No Signal Peptides | 0.944±0.035 | 0.972± 0.018 | 0.954± 0.033 | 0.986± 0.019 | 0.996± 0.005 |

## Appendix B

Hyper-parameters Model Optimization

List of hyper-parameters that were used to optimize the machine learning models during GridSearch.

**Table S2:** kNN Hyperparameters Set During GridSearch. Hyperparameters screened during the k-nearest neighbors (kNN) GridSearch during model training.

| Hyperparameter | Values |
| --- | --- |
| n_neighbors | 1, 3, 5, 7, 9 |
| p | 1, 2, 3 |

**Table S3:** SVC Hyperparameters Set During GridSearch. Hyperparameters screened during the Support Vector Classification (SVC) GridSearch during model training.

| Hyperparameter | Values |
| --- | --- |
| C (Regularization Strength) | 0.1, 0.3, 1, 3, 10, 30 |
| Gamma (Influence) | 0.0001, 0.0003, 0.001, 0.003, 0.01 |

**Table S4:** LR Hyperparameters Set During GridSearch. Hyperparameters screened during the Logistic Regression (LR) GridSearch during model training.

| Hyperparameter | Values |
| --- | --- |
| C (Regularization Strength) | 0.0001, 0.001, 0.01, 0.1, 1, 10 |
| Solver | "liblinear", "sag", "saga", "newton-cg" |
| Penalty | L2 |

**Table S5:** RF Hyperparameters Set During GridSearch. Hyperparameters screened during the Random Forest (RF) GridSearch during model training.

| Hyperparameter | Values |
| --- | --- |
| Max Features | 3, 4, 5, 6 |
| Max Leaf Nodes | 25, 50, 75 |
| n_Estimators (Number of Trees) | 1000 |

**Table S6:** XGB Hyperparameters Set During GridSearch. Hyperparameters screened during XGBoost GridSearch during model training.

| Hyperparameter | Values |
| --- | --- |
| min_child_weight | 3, 4, 5, 6 |
| max_depth (Max Depth of Tree) | 25, 50, 75 |
| gamma (Minimum Loss Reduction to Split) | 0.001, 0.01, 1, 10 |
| reg_lambda (Regularization Term) | 0.001, 0.1, 1, 10 |
| subsample (Percentage of Observations Used Per Tree) | 1 |
| learning_rate (Learning Rate) | 0.05, 0.1, 0.2 |
| Objective (Loss Function) | "binary:logistic" |

## Appendix D

Potential toxin

The regex search applied to the NCBI-derived control set identified over 1,300 proteins with potential toxin activity based on their descriptions. When cross-referenced with UniProt annotations, only 3.34% of these sequences carried the UniProt KW0800 keyword (indicating toxicity). This mismatch between descriptions and annotations highlights inconsistencies in protein labeling within public databases.

**Table S7:**
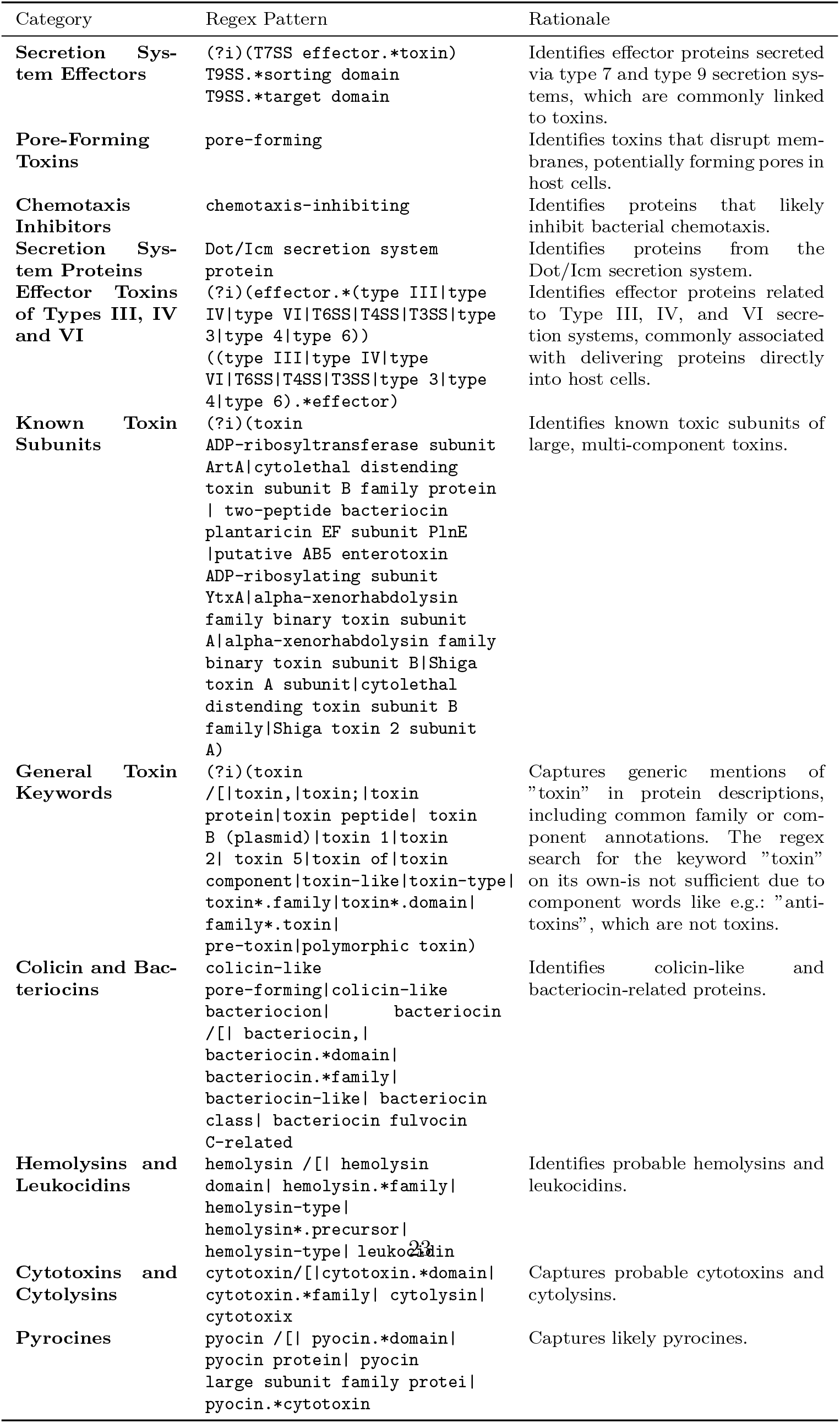
**List of Regex terms part 1** that likely indicate toxicity found in a control set of bacterial proteins.

**Table S8:**
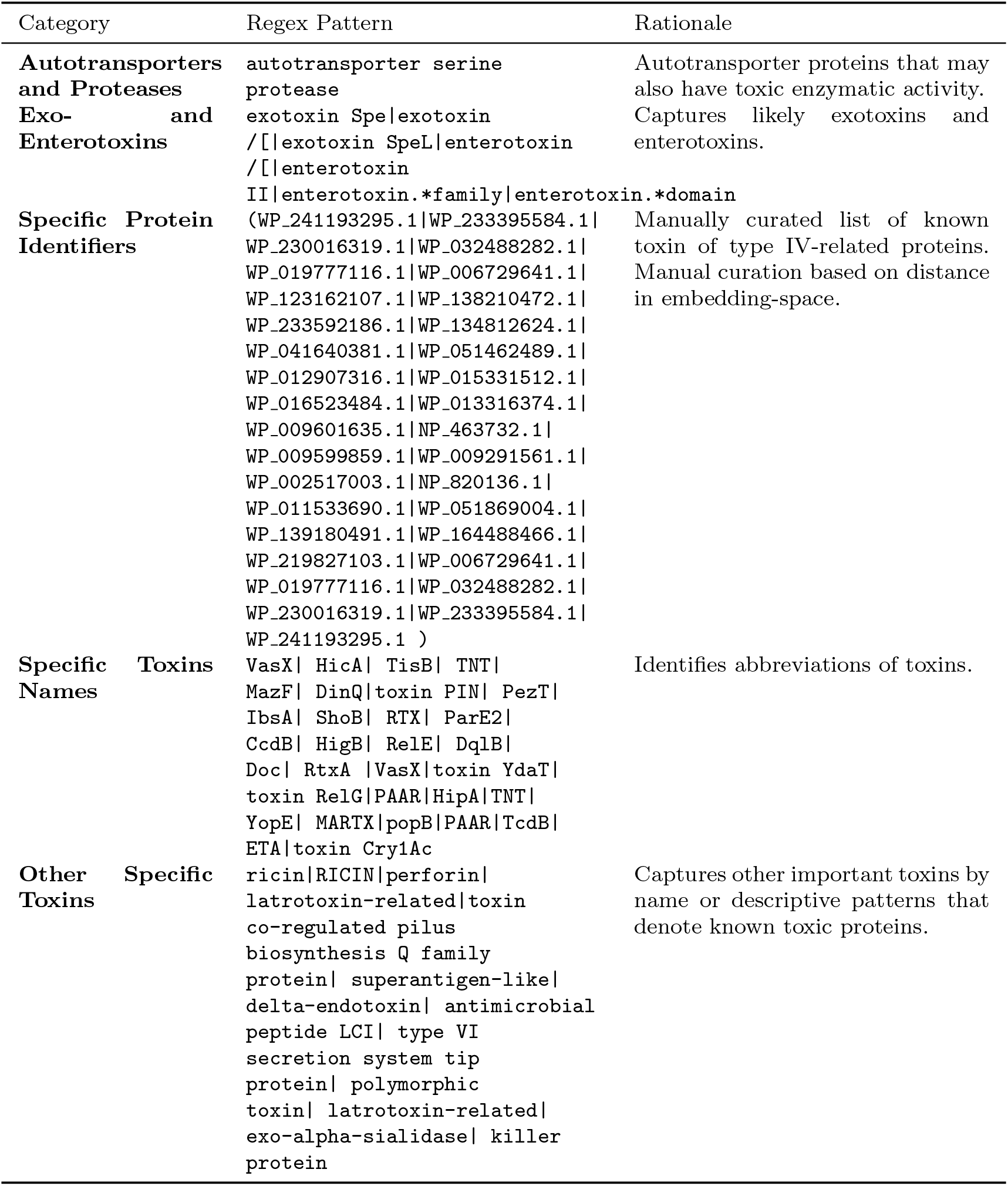
**List of Regex terms part 2** that likely indicate toxicity found in a control set of bacterial proteins.

